# Endothelial SARS-CoV-2 infection is not the underlying cause of COVID19-associated vascular pathology in mice

**DOI:** 10.1101/2023.07.24.550352

**Authors:** Siqi Gao, Alan T. Tang, Min Wang, David W. Buchholz, Brian Imbiakha, Jisheng Yang, Xiaowen Chen, Peter Hewins, Patricia Mericko-Ishizuka, N Adrian Leu, Stephanie Sterling, Avery August, Kellie A. Jurado, Edward E. Morrisey, Hector Aguilar-Carreno, Mark L. Kahn

## Abstract

Endothelial damage and vascular pathology have been recognized as major features of COVID-19 since the beginning of the pandemic. Two main theories regarding how Severe Acute Respiratory Syndrome Coronavirus 2 (SARS-CoV-2) damages endothelial cells and causes vascular pathology have been proposed: direct viral infection of endothelial cells or indirect damage mediated by circulating inflammatory molecules and immune mechanisms. However, these proposed mechanisms remain largely untested in vivo. Here, we utilized a set of new mouse genetic tools^1^ developed in our lab to test both the necessity and sufficiency of endothelial human angiotensin-converting enzyme 2 (hACE2) in COVID19 pathogenesis. Our results demonstrate that endothelial ACE2 and direct infection of vascular endothelial cells does not contribute significantly to the diverse vascular pathology associated with COVID-19.

## Introduction

The most common clinical feature reported in patients with COVID-19 is respiratory symptoms^2,3^. In addition to primarily cause pulmonary symptoms, COVID-19 disease is accompanied by vascular pathology and endothelial damage. Reports emerged around the world confirming a disproportionate prevalence of abnormal thrombotic events and vascular pathology in COVID-19 patients, even in those not in intensive care units^4-11^. Theories regarding the mechanism of vascular pathology observed in COVID-19 patients have been proposed, including direct infection of endothelial cells and systemic inflammatory responses^12-19^. However, these hypotheses remain largely untested and cellular basis of vascular pathology remains controversial. In this study, we used a set of new mouse genetic tools^1^ to rigorously test endothelial contribution to COVID19-associated vascular pathology.

## Results and Discussion

Cellular expression of ACE2 is indispensable for SARS-CoV-2 infection^20,21^, but SARS-CoV-2 is unable to bind mouse ACE2. To determine if endothelial cells directly contribute to lethal infection, we generated animals that express human ACE2 (hACE2) from the mouse Ace2 locus in a manner that enables cell-specific loss of hACE2 using Cre recombinase (*hACE2*^*fl/y*^ mice)^1^. We crossed *hACE2*^*fl/y*^ mice onto a *Tie2-Cre* transgenic mouse line that drives Cre expression in endothelial cells (ECs) to generate mice that express hACE2 in all cells except vascular ECs. *hACE2*^*fl/y*^; *Tie2-Cre*^*+*^ mice and control littermates were exposed to 10^5^ PFU of SARS-CoV-2 virus via nasal inhalation. *hACE2*^*fl/y*^; *Tie2-Cre*^*+*^ mice showed no significant difference in survival after exposure to SARS-CoV-2 compared with the littermate controls (Figure 1A). Histologic analysis revealed the presence of alveolar infiltrates and pulmonary vascular thrombi in the lungs of infected *hACE2*^*fl/y*^; *Tie2-Cre*^*+*^ mice that were indistinguishable from findings observed in control *hACE2*^*fl/y*^ mice (Figure 1B). Expression of inflammation-induced protein intracellular adhesion marker 1 (ICAM1) and the pro-coagulant, inflammation-induced protein von Willebrand’s Factor (vWF) were also similar in the lung capillary endothelial cells of SARS-CoV-2-infected *hACE2*^*fl/y*^ and *hACE2*^*fl/y*^; *Tie2-Cre*^*+*^ mice (Figure 1C and 1D).

**Figure 1.**
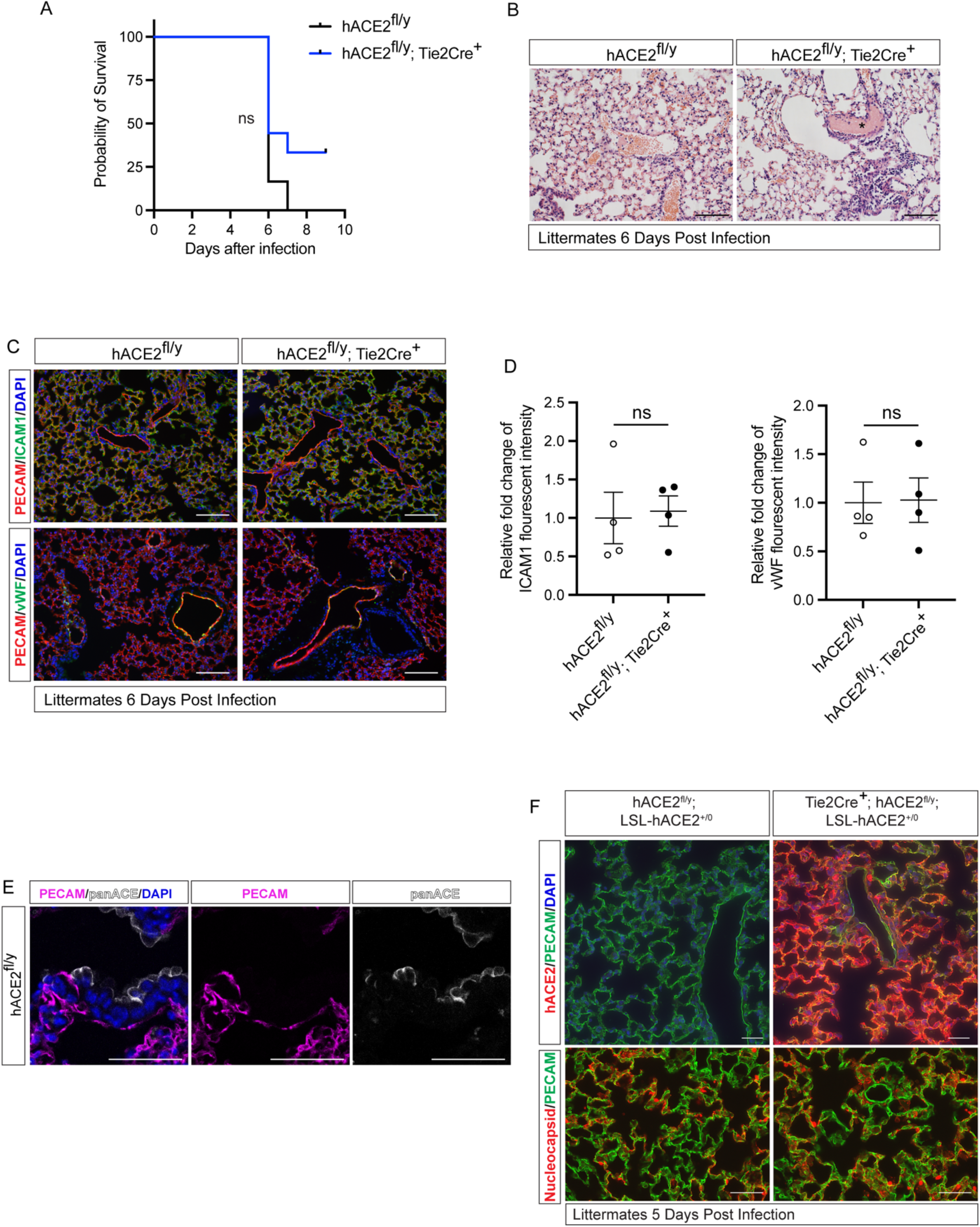
Loss or gain of endothelial hACE2 does not alter SARS-CoV2 infection. **A**, Survival of *hACE2*^*fl/y*^ and *hACE2*^*fl/y*^; *Tie2-Cre*^*+*^ mice (12 to 16-week-old males) after infection with 10^5^ PFU of SARS-CoV-2 via intranasal administration. This viral inoculation method was used in all experiments. n=6 (*hACE2*^*fl/y*^) and 9 (*hACE2*^*fl/y*^; *Tie2-Cre*^*+*^); ns, nonsignificant; data are from two independent experiments. **B**, H&E staining of *hACE2*^*fl/y*^ and *hACE2*^*fl/y*^; *Tie2-Cre*^*+*^ lung tissue 6 days after infection. Asterisk indicates intravascular thrombosis. Scale bars: 100 μm. **C**, Immunofluorescent staining of lung from *hACE2*^*fl/y*^ and *hACE2*^*fl/y*^; *Tie2-Cre*^*+*^ mice with antibodies against ICAM1 or vWF (green), and PECAM (red). Images are representative of N = 4 animals per genotype. Scale bars: 100 μm. **D**, Quantification of ICAM1 and vWF fluorescent intensity. The error bars represent mean±s.d; statical analyses were performed using an unpaired two-tailed t test; ns, nonsignificant. **E**, Immunofluorescent staining of *hACE2*^*fl/y*^ lung tissue using pan-ACE2 antibodies (grey) that recognize both hACE2 and mACE2 proteins and co-stained with PECAM (magenta). Images are representative of N = 3 animals. Scale bars 50 μm. **F**, Immunofluorescent staining of lung from *hACE2*^*fl/y*^; LSL-hACE2^+/0^ and Tie2Cre^+^; *hACE2*^*fl/y*^; LSL-hACE2^+/0^ mice is performed using anti-hACE2 antibody or anti-SARS-CoV-2 nucleocapsid (red) and costained with PECAM (green) 5 days after infection with SARS-CoV-2. The *hACE2*^*fl/y*^ allele enables these mice to be productively infected intranasally. Representative of N = 3 animals per genotype. Scale bars 100 μm.

The studies described above suggested that endothelial cell infection is not required for vascular COVID-19 pathology when hACE2 is expressed at endogenous levels. In fact, immunostaining of lung sections using anti-ACE2 antibodies was able to detect ACE2 expression in epithelial but not endothelial cells^1^ (Figure 1E). To more rigorously test the role of endothelial hACE2 we next crossed *Tie2-Cre* onto a recently described Cre-activated gain of function hACE2 allele (loxP-stop-loxP-hACE2 or LSL-hACE2^+/0^)^1^ to over-express hACE2 in vascular endothelial cells. *Tie2-Cre*;LSL-hACE2^+/0^ animals exhibited very high endothelial-specific expression of hACE2, assessed by immunostaining of tissue sections compared with *hACE2*^*fl/y*^ mice (Figure 1F). To ensure that *Tie2-Cre*;LSL-hACE2^+/0^ animals would be productively infected following SARS-CoV-2 exposure, we generated *Tie2-Cre*;LSL-hACE2^+/0^;*hACE2*^*fl/y*^ animals that support robust infection of the nasal and respiratory epithelium^1^ (Fig. 1F). Despite high levels of endothelial hACE2 expression, we failed to detect nucleocapsid protein that colocalized with PECAM^+^ endothelial cells following nasal SARS-CoV-2 infection (Figure 1F). In contrast, we have previously shown that this gain of function allele is sufficient to drive hACE2 expression and support SARS-CoV-2 infection in both neuronal cells and lung epithelial cells^1^. These studies support the conclusion that SARS-CoV-2 does not confer endothelial cell damage and vascular thrombosis through direction viral infection of those cells. They further demonstrate that the levels of circulating virus are too low to infect even endothelial cells that express very high levels of hACE2, and therefore that most COVID-19 pathology arises due to aerosol infection of the nasal and pulmonary epithelium.

The murine vascular endothelial loss and gain of function studies reported here provide strong in vivo evidence that endothelial ACE2 and direct infection of vascular endothelial cells does not contribute significantly to the diverse vascular pathology associated with COVID-19. These findings are consistent with previously reported in vitro studies that showed human endothelial cells are not readily infected by SARS-CoV-2^18^. Together with our recently reported studies, these findings strongly support a mechanism in which SARS-CoV-2 infection of nasal epithelial and neuronal cells stimulates a powerful inflammatory response that is the cause of COVID-19 vascular pathology.

## Materials and Methods Mice

### Mice

*hACE2*^*fl/y*^ mice, LSL-hACE2^+/0^ mice, and Tie2-Cre transgenic mice have been described^1,22^. All mice were maintained on a mixed genetic background at the University of Pennsylvania animal facility. Mice were genotyped by PCR as described^1^.

### Viral inoculation and tissue harvest

Viral inoculations were performed as described previously^1^. Briefly, mice were anesthetized with isoflurane then intranasally infected with SARS-CoV-2 (Isolate USA-WA1/2020) was obtained from BEI Resource. Mice were monitored and weighed daily, then euthanized at humane endpoint when they lost 20% of the starting weights. Mice studies were combined results from Penn ABSL3 laboratory and Cornell ABSL3 laboratory with accordance to protocols approved by the IACUC at University of Pennsylvanian and Cornell University. For tissue harvest, mice were euthanized with ketamine/xylazine. Lungs were gently inflated with PBS infusion via trachea cannulation. Then lungs were fixed in 4% paraformaldehyde with a minimum of 72 hours to ensure viral inactivation. Tissues were removed from the animal BSL3 facility and followed by ethanol dehydration and embedding in paraffin blocks for histology.

Hematoxylin and eosin staining was performed on paraffin sections.

### Immunofluorescence staining and analysis

Immunohistochemistry staining was performed as previously described^1^ with control and experimental samples on the same slide and under identical staining conditions. Primary antibodies: pan-ACE2 (goat, 1:1,000, R&D AF933), hACE2 (rabbit, 1:200, Abcam ab108209), SARS-CoV-2 nucleocapsid (rabbit, 1:500, Rockland 200-401-A50), ICAM-1 (rabbit, 1:500, Abcam ab179707), vWF (rabbit 1:1,000, Novus Biologicals NB600-586), PECAM (goat 1:500, R&D AF3628). Fluorescence-conjugated Alexa Fluor secondary antibodies were used (1:500, Invitrogen) according to the primary antibody species and counterstained with DAPI (1:1000). ICAM1and vWF fluorescence intensity was calculated by Integrated fluorescence intensity. All images were analyzed using ImageJ/FIJI software.

### Statistics

Mice were inoculated with SARS-CoV-2 in blinded fashion without knowledge of genotypes, and infections were performed at two separate ABSL-3 facilities with independent experimenters. Statistical tests used to determine significance are described in the figure legends. Graph generation and statistical analyses performed with GraphPad Prism 9.5.1. Survival curve statistics were performed with log-rank Mantel Cox tests. All *t* tests performed were two-tailed.

## Acknowledgments

We thank Kahn lab members for helpful discussions.

